# Differential expression and modulation of presenilin-1 and presenilin-2 in neural cell lines

**DOI:** 10.1101/2024.10.15.618376

**Authors:** Melissa K. Eccles, Amy Woodfield, Ayeisha Milligan Armstrong, Mark Agostino, David M. Groth, Giuseppe Verdile

**Affiliations:** Centre for Genomic & Experimental Medicine, Institute of Genetics & Cancer, University of Edinburgh, Western General Hospital, Edinburgh, UK; Curtin Medical School, Curtin Health Innovation Research Institute (CHIRI), Curtin University, Bentley, Western Australia, Australia; Curtin Institute for Computation, Curtin University, Bentley, Western Australia, Australia; School of Medical and Health Sciences, Edith Cowan University, Joondalup, Western Australia, Australia

**Author notes:** Correspondence to: **Melissa K. Eccles**, Centre for Genomic & Experimental Medicine, Institute of Genetics & Cancer, University of Edinburgh, Western General Hospital, Crewe Rd, Edinburgh EH4 2XU, UK. Funding Information: MKE - Dementia Australia | Dementia Australia Research Foundation (DARF).

**Keywords:** Presenilin-1, Presenilin-2, Alzheimer’s disease, neurons, microglia

## Abstract

Presenilin proteins, the catalytic components of the γ-secretase enzyme, play a key role in the pathogenesis of Alzheimer’s disease primarily by generating amyloid-β peptides. Although often grouped together, Presenilin-1 (PS1) and Presenilin-2 (PS2) are distinct proteins that contribute differently to γ-secretase activity. This study examines the differential expression of PS1 and PS2 in various neuroblastoma and microglial cell lines, along with compensatory responses following the ablation of one presenilin homologue. Using quantitative immunoblotting, we show that PS1 and PS2 expression levels vary significantly across cell types. Notably, the ablation of PS2 results in increased PS1 expression, particularly in microglial cells, highlighting the importance of PS2. Additionally, neuronal differentiation of two neuroblastoma lines, caused changes in protein expression levels resulting in similar expression profiles of PS1 and PS2, with PS2 levels being higher than PS1. Understanding PS expression profiles is crucial for distinguishing between PS1- and PS2-mediated γ-secretase activity and for developing γ-secretase-targeted therapeutics with improved selectivity.

## INTRODUCTION

The enzyme γ-secretase cleaves numerous substrates essential for cellular function and plays a crucial role in Alzheimer’s disease (AD) by generating amyloid-β (Aβ) peptides (Kimberly et al., 2000). In AD, an elevated Aβ42:40 ratio is often observed, indicating a relative increase in the production of the more aggregation-prone Aβ42 peptide compared to Aβ40, which accelerates amyloid plaque formation, a key hallmark of the disease (Gouras et al., 2000). Mutations in the catalytic component, presenilin, are known to cause autosomal dominant forms of AD (Ryman et al., 2014), typically increasing the Aβ42:40 ratio (Tang and Kepp, 2018). While often referred to in the literature as a singular “presenilin” protein (Inuzuka et al., 2011; Muller et al., 2023) there are two distinct proteins – Presenilin-1 (PS1) and Presenilin-2 (PS2) – both of which can form γ-secretase enzymes (Yonemura et al., 2016), herein referred to as PS1γ and PS2γ; these enzymes exhibit distinct substrate specificities and localisations (Meckler and Checler, 2016; Sannerud et al., 2016; Watanabe et al., 2021). A critical role for PS1 in murine embryogenesis has been well established (Donoviel et al., 1999; Herreman et al., 1999; Mastrangelo et al., 2005; Shen et al., 1997; Wong et al., 1997). However, compared to younger mice the ratio of PS1 to PS2 expression decreases in aged mouse brains (Ghosh and Thakur, 2008; Kaja et al., 2015; Placanica et al., 2009b; Thakur and Ghosh, 2007). These findings align with human transcript data, suggesting that, in the cortex, *PSEN2* expression increases with age such that the expression of *PSEN2* is higher than *PSEN1*, unlike the foetal cortex where *PSEN1* expression is higher (Lee et al., 1996). This increased PS2 expression with age may be significant in the context of AD as it has been shown that PS2γ generates higher Aβ42:Aβ40 than PS1γ (Eccles et al., 2024; Pimenova and Goate, 2020; Placanica et al., 2009a; Watanabe et al., 2021).

Our understanding of the functional roles of presenilin, like many other proteins, has been shaped by the use of knockout mouse models, cell lines, and the subsequent rescue in in vitro cell lines. However, the presence of two homologous presenilin proteins, with structural and functional overlap, complicates the interpretation of results. A large portion of studies fail to acknowledge the presence of two presenilin proteins, often knocking out PS1 and assigning the observed outcome to PS1 with no consideration of PS2. Is the observed loss/gain of function a result of the loss of the ablated homologue or the possible upregulation of the alternate homologue? In vitro studies utilising the expression of ectopic PS1 or PS2 on a PSnull background rarely tag PS proteins, with only a few studies having tagged the exogenous PS and compared corresponding expression levels (Eccles et al., 2024; Sannerud et al., 2016; Yonemura et al., 2011). This is likely because it is assumed that ectopic expression, using constitutive promoters, generates similar expression levels of closely homologous proteins (Xia et al., 2006). We have shown that this assumption is incorrect for overexpressed PS1 and PS2 incorporation into γ-secretase (Eccles et al., 2024), which is likely due to regulation of the γ-secretase complex formation by the other critical components of the enzyme (Edbauer et al., 2002; Kimberly et al., 2003; Luo et al., 2003; Steiner et al., 2002; Thinakaran et al., 1997; Yang et al., 2002). In this study we demonstrate the importance of understanding PS protein expression when interpreting functional consequences across different cell types, using our recently developed tool that enables absolute quantitation of endogenous PS1 and PS2 (Eccles et al., 2024).

## RESULTS AND DISCUSSION

PS1 and PS2 levels were quantitatively assessed via immunoblot using the PS-Std and method previously described (Eccles et al., 2024), in human neuroblastoma BE(2)-M17 (herein referred to as M17) and SH-SY5Y cells, and microglial HMC3 cells (Fig 1). SH-SY5Y cells genetically modified to overexpress APP695 or APP695swe are also commonly used, and so were also assessed. This method determines the absolute quantitation of PS1 and PS2 expression allowing for direct comparison of the PS1 and PS2 expression data across the multiple experiments. Comparison of the PS1 and PS2 expression levels in cells with the PS1+/+PS2+/+ (WT) genotype show that PS1 expression is significantly higher than PS2 expression in all cell lines except the M17 cell line, where PS2 expression trends toward being higher than PS1 expression (p=0.0508; Fig. 2A). The PS expression profile, determined as the PS1:PS2 ratio for all cell lines, was compared to all other cell lines (Fig. 2B). The PS1:PS2 ratios demonstrate a very broad range from over 20-fold greater PS1 than PS2 in HMC3 cells to almost 2-fold greater PS2 than PS1 in M17 cells, and were significantly different between cells of different origin. Only the SHSY5Y cell lines showed no significant difference in PS1:PS2, indicating that the over-expression of the APP substrate in these cells does not alter the PS expression profile. Overall, these findings underscore the need to determine comparative PS expression levels in cellular models of AD. It emphasizes the importance of assessing protein expression in the specific cell line of interest before making assumptions about the effects of protein ablation on the functional phenotype.

**Figure 1:**
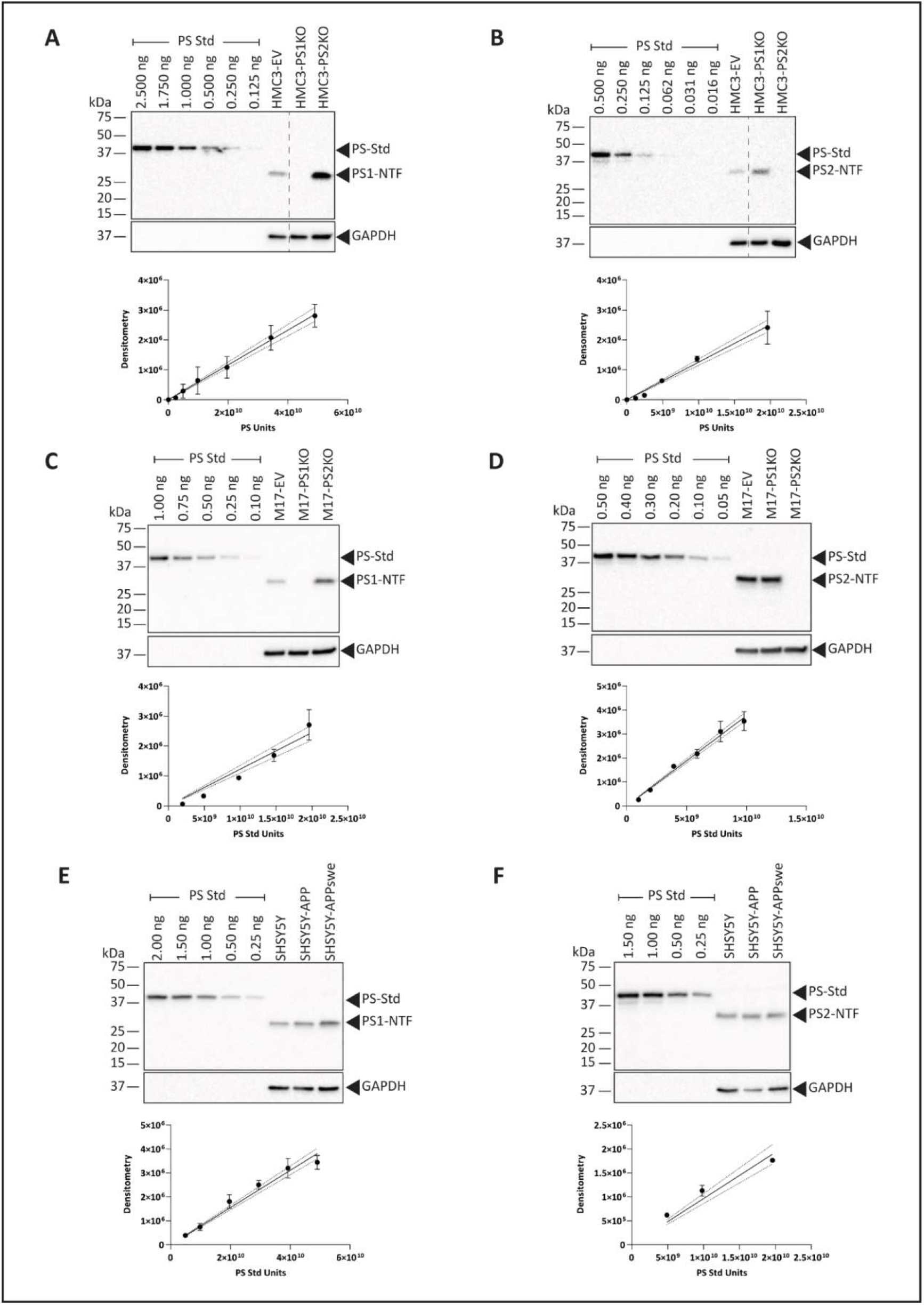
PS1-NTF and PS2-NTF expression in HMC3, M17 and SHSY5Y cells. Immunoblot results and PS-Std standard curves for PS1-NTF (A, C, E) and PS2-NTF (B, D, F) for HMC3-EV, -PS1KO and -PS2KO cell lines (A-B), M17-EV, -PS1KO, and -PS2KO cell lines (C-D), and SH-SY5Y, SH-SY5Y-APP, and SH-SY5Y-APPswe cell lines (E-F). Representative blot of 3 to 4 replicates presented.

**Figure 2:**
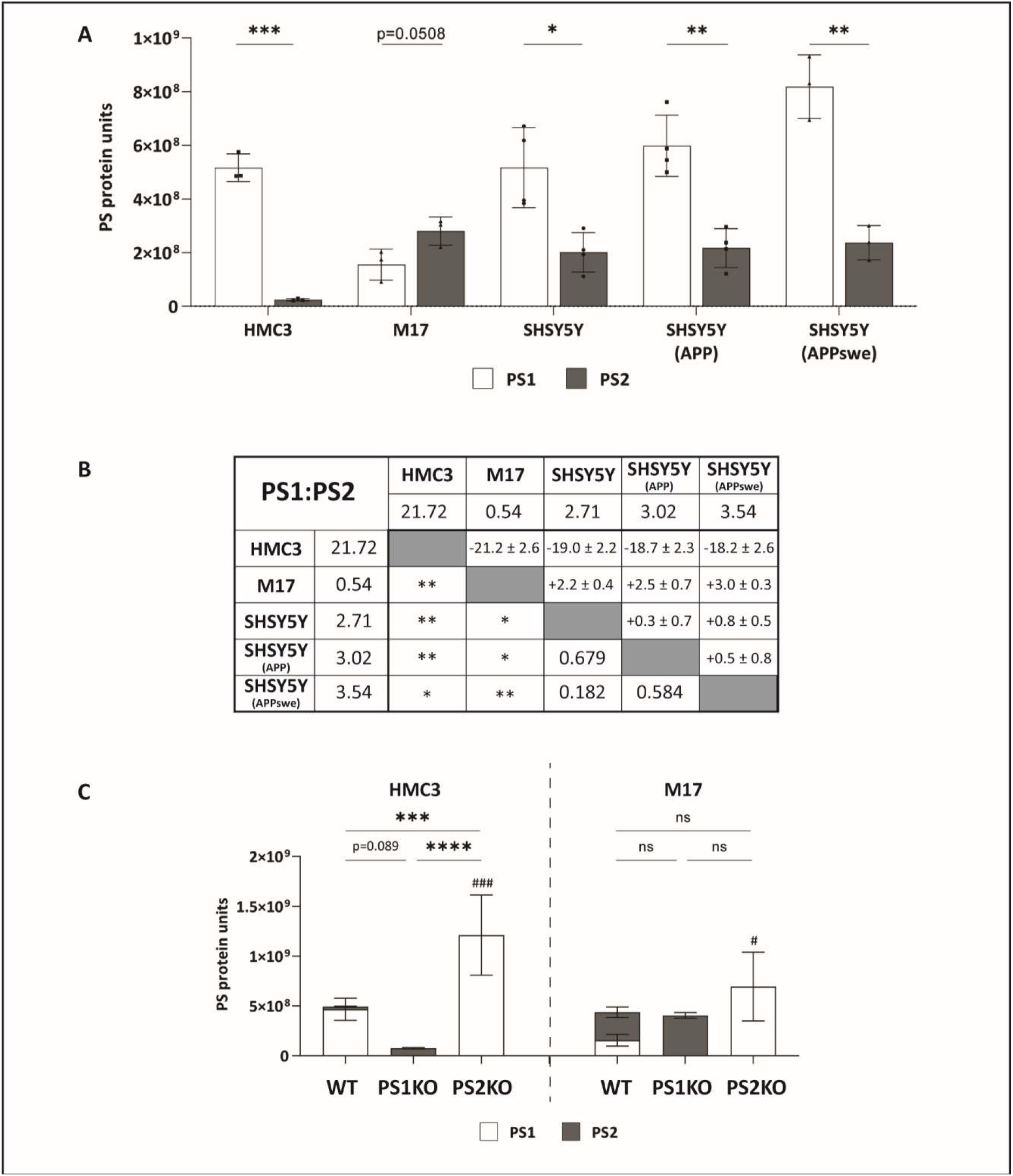
PS protein unit analysis in WT and PS deficient cell lines. Quantiation of PS protein units in cell lines with WT genotype (A). PS1:PS2 ratio comparison matrix between cell lines where data above the diagonal is the difference in the PS1:PS2 ratio between the two cell lines as determined by unpaired t-test, below the diagonal is the statistical significance of the unpaired t-test after FDR correction for multiple testing (B). PS protein units in HMC3 and M17 cells with altered PS genotypes, significant statistical differences in PS1 expression represented by #, no differences in PS2 expression reach statistical significance, results of statistical assessment of the sum of PS1 and PS2 represented by * or ns (C). Values shown are mean ± SD. Statistical tests applied were unpaired t-test, with FDR correction for multiple testing (A, B), two-way ANOVA with Holm-Šidák’s multiple comparison test (within cell lines) (C) where */# = P < 0.05, **/## = P < 0.01, ***/### = P < 0.001, ****/#### = P <0.0001.

The loss of PS1 has previously been shown to result in a compensatory, relative, upregulation of PS2 expression, and vice-versa in HEK293T cells (Lessard et al., 2019) and HEK293 cells (Eccles et al., 2024); as such, we sought to investigate this response in the HMC3 and M17 cells lines. To achieve this, PS1 or PS2 were ablated from HMC3 and M17 cells using CRISPR-Cas9. Three clones each of genotype PS1-/-PS2+/+ (PS1KO) and genotype PS1+/+PS2-/- (PS2KO) cell lines for both HMC3 and M17, as well as empty vector controls, were generated. These clones were assessed via immunoblotting and qPCR to identify a representative clone for use in subsequent experiments. (SI Fig 1; SI Fig 2).

To demonstrate the magnitude of the changes in total PS and absolute PS1:PS2 expression levels, PS1 and PS2 protein units are presented as stacked graphs in Figure 2C. The HMC3 WT cells expressed the lowest PS2 levels, compared with other cells of the same genotype. However, when PS2 was ablated in these cells, PS1 expression increased significantly by 2.6-fold. In contrast, ablation of PS1 did not lead to a significant change in PS2 levels. Consequently, the loss of PS2 leads to significantly more total PS protein, while the loss of PS1 results in a trend (p=0.089) toward lower total PS protein. These data indicate that PS2 could be more critical than PS2 in microglial cells, which supports previous work showing PS2 as the predominant presenilin in murine microglia (Jayadev et al., 2010).

Similarly, in M17 cells, loss of PS2 led to an increase in PS1 expression, while the loss of PS1 had no effect on PS2 expression. However, as the initial level of PS2 was higher than PS1 in the WT cells, no difference between the total PS units expressed with the loss of either PS1 or PS2 was observed. This suggests that, in the M17 neuroblastoma cells, PS1 and PS2 may have considerable functional overlap where the loss of either homologue results in compensatory expression resulting in overall equal total PS expression. Interestingly, while HMC3 and M17 cell lines have comparable levels of total PS protein expression, they have vastly different PS1:PS2 profiles; both demonstrate a compensatory increase in PS1 with the loss of PS2, but no significant effect on PS2 expression with the loss of PS1. This is in contrast to previous reports in HEK293-derived cell lines that demonstrate that the alternate homologue compensates for the loss of either PS1 or PS2 (Eccles et al., 2024; Lessard et al., 2019).

It has previously been reported that PS1 (Flood et al., 2004) and PS2 (Culvenor et al., 2000; Watanabe et al., 2021) expression can increases in cells with neuronal differentiation. Here, PS1 and PS2 expression levels were quantitated in differentiated M17 and SH-SY5Y cell lines to directly compare PS expression in response to neuronal differentiation. Cells were cultured for 7 days with either 10 mM retinoic acid (M17 cells) or 10 nM staurosporine (SH-SY5Y cells) (Andres et al., 2013; Filograna et al., 2015), to induce neuronal differentiation, or with vehicle control. Whole cell lysates were subsequently immunoblotted for detection of PS1 and PS2 (Fig 3A-B), alongside the PS-Std, to quantitate and compare PS1 and PS2 protein expression.

**Figure Error! No text of specified style in document.3:**
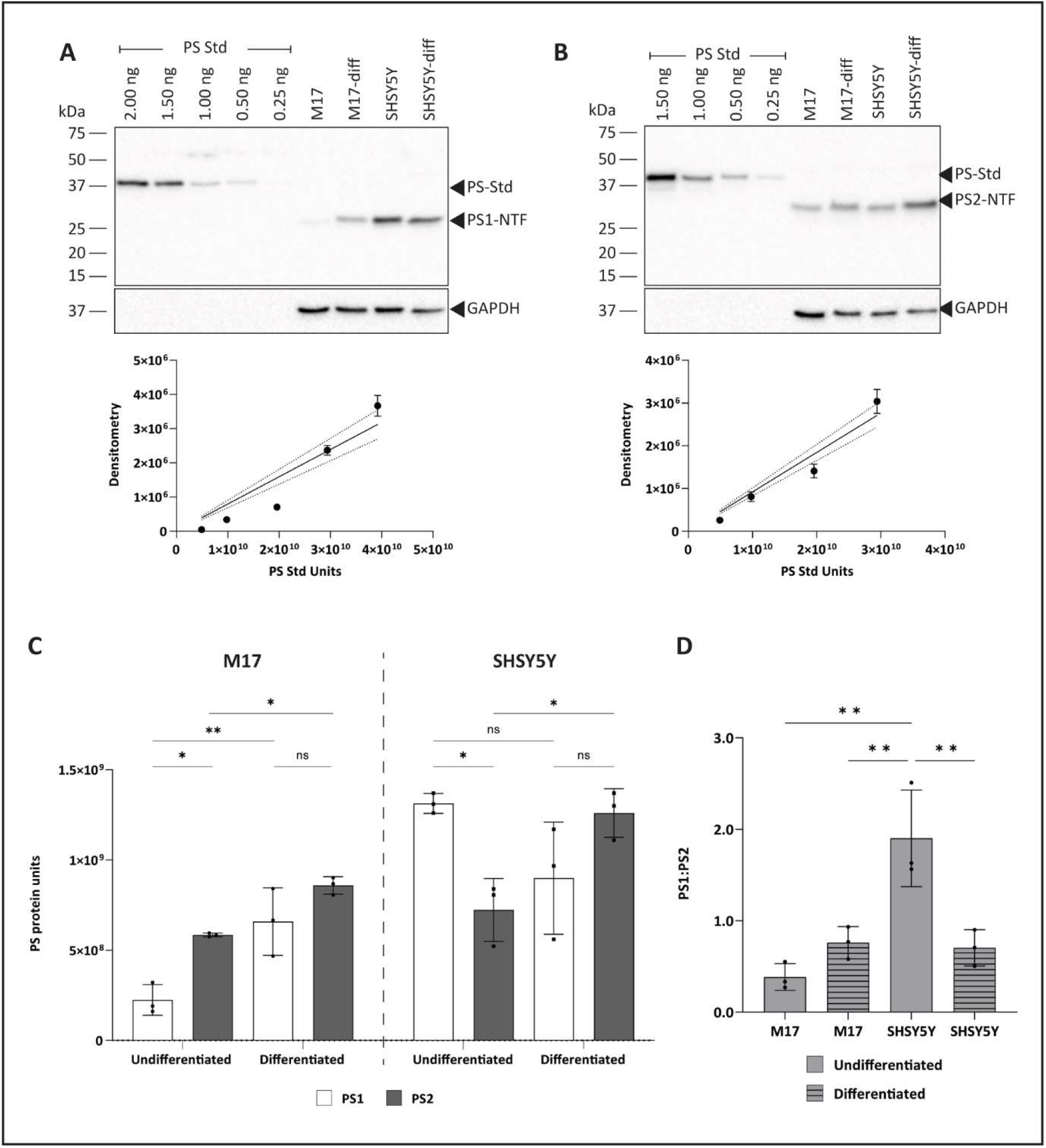
Effect of differentiation on PS expression in neuroblastomas. Immunoblot results and PS-Std curves of PS1-NTF (A) and PS2-NTF (B) in M17 and SH-SY5Y neuroblastoma cell lines with and without differentiation, representative blots of 3 biological replicates. Quantiated PS protein units in differentiated and undifferntiated M17 and SH-SY5Y cells (C). Ratio of PS1:PS2 protein levels in differentiated and undifferntiated M17 and SH-SY5Y cells (D). Values shown are mean ± SD. Statistical tests applied were two-way ANOVA (C) and one-way ANOVA (D) with Holm-Šidák’s multiple comparison test where ns = not significant, * = P < 0.05, ** = P < 0.01, *** = P < 0.001, **** = P <0.0001

In undifferentiated cells (Fig 3C), the level of PS1 expression was significantly lower than PS2 expression in the M17 cells. Conversely, in SH-SY5Y cells PS1 protein expression was significantly higher than PS2. This difference is also reflected in the markedly higher in PS1:PS2 ratio observed in SH-SY5Y cells compared to M17 cells (Fig 3D). Interestingly, in both M17 and SH-SY5Y cells, PS2 expression responded similarly, increasing in expression with differentiation (Fig 3C). However, the effect of differentiation on PS1 expression was discordant between the two cell lines; an increase was observed in M17 cells, while no change occurred in SH-SY5Y cells (Fig 3C). Consequently, the resulting PS1:PS2 ratio is the same in both differentiated cell lines (M17 = 0.76; SH-SY5Y = 0.70) (Fig 3D). These findings demonstrate that the PS expression profiles of M17 and SH-SY5Y differ significantly in the undifferentiated sate. The higher PS1 protein levels in the SH-SY5Y cells may be a function of elevated Notch1 protein levels compared to M17 (Ross et al., 2015). Notch1 localises to the plasma membrane, the predominant site of PS1γ localisation (Sannerud et al., 2016), where it undergoes canonical Notch signalling and processing by γ-secretase (Kopan and Ilagan, 2009). Upon differentiation to a more neuronal-like state, the PS1:PS2 ratio becomes similar, with both cell lines showing higher PS2 expression than PS1 expression. Interestingly, Notch1 expression decreases with neuronal differentiation (Ross et al., 2015).

Knowledge of PS expression levels is necessary for ascertaining the specific contributions of PS1γ versus PS2γ. It has been observed that the Aβ42:Aβ40 ratio increases, while the PS1:PS2 expression ratio decreases, with age in murine models (Placanica et al., 2009b). Similarly in mice, where PS1 is conditionally ablated, but PS2 expression is retained, the Aβ42:Aβ40 ratio is increased (Yu et al., 2001). The observed increase in Aβ42:Aβ40 ratio may be a consequence of decreased PS1:PS2 expression ratio, particularly with age, as several studies have shown that PS2γ generates higher Aβ42:Aβ40 ratio than PS1γ (Eccles et al., 2024; Pimenova and Goate, 2020; Placanica et al., 2009a; Watanabe et al., 2021). Advances in single cell RNA sequencing technologies are providing new insights into *PSEN1* and *PSEN2* expression in neurons and glial cells, where *PSEN2* is more highly expressed in neurons, compared with *PSEN1* which is mostly highly expressed in oligodendrocytes (Herring et al., 2022; Kelley et al., 2018). The current study presents two key results that suggest PS2 may play a more important role in neurons. Firstly, the loss of either PS1 or PS2 in M17 cells results in compensatory upregulation of the alternate PS homologue, such that there is no significant difference in total PS expression. Secondly, PS2 upregulation occurs in both M17 and SH-SY5Y cells that have been differentiated, to be more neuronally representative, such that the PS1:PS2 ratios are <1.0. These results suggest important roles for both PS1 and PS2 in mature neurons, and support previous results in induced pluripotent stem cells showing an increase of PS2 expression with differentiation, concomitant with no change in PS1 expression (Watanabe et al., 2021).

Further to the AD-associated functional roles of PS1 and PS2 in neurons (Cao et al., 2018; Feng et al., 2004; Kang and Shen, 2020; Lauer et al., 2020; Peng et al., 2022; Saura et al., 2004; Soto-Faguás et al., 2021; Watanabe et al., 2021; Watanabe et al., 2014; Wines-Samuelson et al., 2010; Zhu et al., 2008), important roles for PS in microglial cytokine response (Jayadev et al., 2013; Jayadev et al., 2010) and Aβ clearance (Farfara et al., 2011; Wunderlich et al., 2013) have also been reported. In addition, several proteins important for the microglial response to Aβ are substrates of γ-secretase (Fourgeaud et al., 2016; Güner and Lichtenthaler, 2020; Huang et al., 2021; Kemmerling et al., 2017; Kim et al., 2009; Manocha et al., 2016; Rangaraju et al., 2018; Wunderlich et al., 2013; Zhao et al., 2020), and PS2 is suggested to be critical for microglial cytokine responses (Jayadev et al., 2013; Jayadev et al., 2010). In contrast to the suggested importance of PS2 in microglial function, it has been observed in this study that of all the cells examined, PS2 expression is lowest in HMC3 cells – reduced by at least an order of magnitude compared to the next lowest SH-SY5Y cells. Interestingly though the loss of this comparatively small amount of PS2 causes a significant increase in PS1 protein levels such that the total PS expression in the PS1KO HMC3 cells is significantly higher than the WT HMC3 cells. This observation aligns with previous reports of significantly increased PS1 expression with loss of PS2 in murine microglia (Jayadev et al., 2010). Although not conclusive evidence, the level of increase of PS1 expression in response to the loss of PS2 is certainly suggestive of a critical role for PS2 in microglial function.

In conclusion, the utility of our PS-Std tool for direct quantitative assessment of endogenous PS1 and PS2 expression has been demonstrated, highlighting the potential applications and novel insights that can be achieved by quantitatively comparing PS protein expression. Recognising that PS1 and PS2 expression is highly variable in different cell types commonly used in the field emphasises the need for due consideration of expression to ensure accurate interpretation of the functional roles of PS1 and PS2 in different settings. This information is essential when investigating the function of these proteins and to enable improved development of selective γ-secretase targeting therapeutics.

## MATERIAL AND METHODS

### CRISPR-Cas9 Presenilin Knock-Out

PS1 and PS2 knockout cell lines were generated in M17 (RRID:CVCL_0167) and HMC3 (Gifted by Ryan Anderton, Perron Institute, RRID:CVCL_II76) cells using pSp-Cas9-(BB)-2A-GFP vector (RRID:Addgene_48138) (Ran et al., 2013). M17 and HMC3 cells were transfected with pSp-Cas9-(BB)-2A-GFP vectors with gRNA sequence targeting either PSEN1, PSEN2, or empty vector for control cell lines. Two sites per gene were targeted simultaneously using a 1:1 ratio of the vectors (SI Table 1), transfected using Lipofectamine 3000 (Invitrogen L3000015) as per manufacturer’s instructions. Twenty-four hours post transfection, cells were trypsinised, prepared in single cell suspension and single cell sorted for mid-level GFP expression as marker for pSp-Cas9-(BB)-2A-GFP transfection, and propidium iodide for live/dead cell differentiation.

### Quantitative PCR

RNA was extracted from cultures at approximately 90% confluence, using ISOLATE II RNA extraction kit as per instructions (Bioline BIO-52072). cDNA was generated using Tetro cDNA kit (Bioline BIO-65043), with the following adaptions: 1) Random hexamers and oligo dTs were used in 1:1 ratio such that final volume of primers was as per recommended amount for reaction volume. 2) Reaction was incubated at room temperature for 10 min followed by 45 °C for 1 hour.

cDNA generated from wild type cell RNA extracts was used to generate standard curves for all qPCR targets to determine appropriate amount of RNA (µg/µl) to be used for specific targets, and primer efficiencies for use in relative quantitation calculations (Pfaffl, 2001). Primer sequences were designed at NCBI PrimerBlast, unless otherwise stated, and are presented in SI Table 2. Quantitative PCR reactions with final primer concentration of 500 nM were prepared using GoTaq qPCR master mix (Promega A6001) in 20 µl reactions and run on a Viia-7 (Invitrogen).

### Neuroblastoma Differentiation

Cells were differentiated as described previously (Andres et al., 2013; Filograna et al., 2015). Briefly, M17 cells were plated at 2187.5 cells/cm^2^, in 100mm dishes. 24 hrs after plating, 10 mM retinoic acid differentiation media was added, and replaced every 24 hrs. SH-SY5Y (RRID:CVCL_0019) cells were plated at 2187.5 cells/cm^2^, in 100mm dishes. 24 hrs after plating, 10 nM staurosporine differentiation media was added, and replaced every 48hrs. Control cells were treated with DMSO. Cells were harvested for lysates after 7 days of differentiation.

### Western Immunoblotting

Protein lysates and PS-Std were prepared for separation by PAGE as previously described (Eccles et al., 2024). Proteins were resolved on 12% w/v tris-tricine poly acrylamide gels by electrophoresis at 100 V for approximately 1 hr 45 min, in tris-tricine cathode (100 mM tris, 100 mM tricine, pH8.3) and anode (200 mM tris, pH8.8) buffer. The resolved proteins were then transferred to 0.2 µm nitrocellulose membrane (BioRad) via wet transfer in 192 mM tris, 25 mM glycine, 20 % v/v methanol buffer, at 150 mA overnight at 4 °C. Membranes were blocked with 5% w/v non-fat dried milk powder (NFDM) in TBS for 1 hr at room temperature, then incubated overnight at 4 °C with either α-PS1 NTF antibody (1:2000; Biolegend PS1 NT1 823401) or α-PS2 NTF antibody (1:1000, Biolegend PS2 814204) prepared in 0.5% w/v skim milk in TBST (TBS with addition of 0.05% v/v Tween-20). Membranes were subsequently washed, followed by incubation with α-mouse IgG HRP conjugated secondary antibody (1:20,000; Thermo Fisher 31430), for 1 hr at room temperature. Membranes were again washed and prepared for imaging by incubating them with either Clarity ECL (BioRad 1705061), for PS1 NTF detection, or Prime ECL (Cytiva GERPN2232), for PS2 NTF detection. Western blots were imaged on BioRad ChemiDoc MP system and band densitometry quantitated using ImageLab (BioRad, version 6.1.0 build 7).

### Endogenous Presenilin Quantitation

Endogenous presenilin expression was quantitated as previously described (Eccles et al., 2024). Briefly, using the PS-Std, an appropriate range was determined via western blotting of representative samples such that the samples would fall within the range of the standard curve. A minimum of four quantitation standards were used to generate the standard curve, and a minimum of 3 replicates of the standard curve were generated concurrently via PAGE and western blot for each experiment. After western blotting with the appropriate PS antibody, the standard curve was quantitated using ImageLab to determine the densitometry for the standard curve. The densitometry (arbitrary units) was plotted against the number of PS-Std units, determined by PS-Std ng / 5.10×10^−11^ ng, where 5.10×10^−11^ ng is the mass of 1 unit of PS-Std. The equation for the standard curve was then used to calculate the number of PS1 or PS2 protein units from the western blot band densitometry, and subsequently normalised per µg of total protein and by loading control.

### Statistical Analysis

All statistical analysis was completed using GraphPad Prism 9.5.0. Three to four biological replicates were completed for all in vitro experiments. Shapiro-Wilk normality test was completed to determine if data was normally distributed. Statistical significance for normally distributed data was determined via unpaired T-test where only two groups were examined. Where more than two groups are examined, one-way ANOVA or two-way ANOVA analysis with Holm-Šidák’s multiple comparison tests are used as appropriate.

## Supporting information

Eccles - Differential expression and modulation of presenilin-1 and presenilin-2 in neural cell lines - SI Material

## Data Availability

The data underlying all figures are available in the published article and its online supplemental material.

## Abbreviations

Aβ: amyloid-β
AD: Alzheimer’s disease
CNS: central nervous system
PS-Std: presenilin fusion standard
PS1: presenilin-1
PS2: presenilin-2
PS1γ: PS1-γ-secretase
PS2γ: PS2-γ-secretase

## Notes

### Competing Interest Statement

The authors have declared no competing interest.

## REFERENCES

Andres, D., B.M. Keyser, J. Petrali, B. Benton, K.S. Hubbard, P.M. McNutt, and R. Ray. 2013. Morphological and functional differentiation in BE(2)-M17 human neuroblastoma cells by treatment with Trans-retinoic acid. BMC neuroscience. 14:49.

Cao, T., X. Zhou, X. Zheng, Y. Cui, J.Z. Tsien, C. Li, and H. Wang. 2018. Histone Deacetylase Inhibitor Alleviates the Neurodegenerative Phenotypes and Histone Dysregulation in Presenilins-Deficient Mice. Front Aging Neurosci. 10.

Culvenor, J.G., G. Evin, M.A. Cooney, H. Wardan, R.A. Sharples, F. Maher, G. Reed, A. Diehlmann, A. Weidemann, K. Beyreuther, and C.L. Masters. 2000. Presenilin 2 expression in neuronal cells: induction during differentiation of embryonic carcinoma cells. Experimental cell research. 255:192–206.

Donoviel, D.B., A.-K. Hadjantonakis, M. Ikeda, H. Zheng, P.S.G. Hyslop, and A. Bernstein. 1999. Mice lacking both presenilin genes exhibit early embryonic patterning defects. Genes & development. 13:2801–2810.

Eccles, M.K., N. Main, R. Carlessi, A.M. Armstrong, M. Sabale, B. Roberts-Mok, J.E.E. Tirnitz-Parker, M. Agostino, D. Groth, P.E. Fraser, and G. Verdile. 2024. Quantitative comparison of presenilin protein expression reveals greater activity of PS2-γ-secretase. The FASEB Journal. 38:e23396.

Edbauer, D., E. Winkler, C. Haass, and H. Steiner. 2002. Presenilin and nicastrin regulate each other and determine amyloid β-peptide production via complex formation. Proceedings of the National Academy of Sciences of the United States of America. 99:8666–8671.

Farfara, D., D. Trudler, N. Segev-Amzaleg, R. Galron, R. Stein, and D. Frenkel. 2011. γ-Secretase component presenilin is important for microglia β-amyloid clearance. Annals of neurology. 69:170–180.

Feng, R., H. Wang, J. Wang, D. Shrom, X. Zeng, and J.Z. Tsien. 2004. Forebrain degeneration and ventricle enlargement caused by double knockout of Alzheimer’s presenilin-1 and presenilin-2. Proceedings of the National Academy of Sciences of the United States of America. 101:8162–8167.

Filograna, R., L. Civiero, V. Ferrari, G. Codolo, E. Greggio, L. Bubacco, M. Beltramini, and M. Bisaglia. 2015. Analysis of the Catecholaminergic Phenotype in Human SH-SY5Y and BE(2)-M17 Neuroblastoma Cell Lines upon Differentiation. PloS one. 10:e0136769–e0136769.

Flood, F., E. Sundström, E.-B. Samuelsson, B. Wiehager, Å. Seiger, J.A. Johnston, and R.F. Cowburn. 2004. Presenilin expression during induced differentiation of the human neuroblastoma SH-SY5Y cell line. Neurochemistry International. 44:487–496.

Fourgeaud, L., P.G. Través, Y. Tufail, H. Leal-Bailey, E.D. Lew, P.G. Burrola, P. Callaway, A. Zagórska, C.V. Rothlin, A. Nimmerjahn, and G. Lemke. 2016. TAM receptors regulate multiple features of microglial physiology. Nature. 532:240–244.

Ghosh, S., and M.K. Thakur. 2008. PS2 protein expression is upregulated by sex steroids in the cerebral cortex of aging mice. Neurochemistry International. 52:363–367.

Gouras, G.K., J. Tsai, J. Naslund, B. Vincent, M. Edgar, F. Checler, J.P. Greenfield, V. Haroutunian, J.D. Buxbaum, H. Xu, P. Greengard, and N.R. Relkin. 2000. Intraneuronal Abeta42 accumulation in human brain. The American journal of pathology. 156:15–20.

Güner, G., and S.F. Lichtenthaler. 2020. The substrate repertoire of γ-secretase/presenilin. Seminars in cell & developmental biology. 105:27–42.

Herreman, A., D. Hartmann, W. Annaert, P. Saftig, K. Craessaerts, L. Serneels, L. Umans, V. Schrijvers, F. Checler, H. Vanderstichele, V. Baekelandt, R. Dressel, P. Cupers, D. Huylebroeck, A. Zwijsen, F. Van Leuven, and B. De Strooper. 1999. Presenilin 2 Deficiency Causes a Mild Pulmonary Phenotype and no Changes in Amyloid Precursor Protein Processing but Enhances the Embryonic Lethal Phenotype of Presenilin 1 Deficiency. Proceedings of the National Academy of Sciences of the United States of America. 96:11872–11877.

Herring, C.A., R.K. Simmons, S. Freytag, D. Poppe, J.J.D. Moffet, J. Pflueger, S. Buckberry, D.B. Vargas-Landin, O. Clément, E.G. Echeverría, G.J. Sutton, A. Alvarez-Franco, R. Hou, C. Pflueger, K. McDonald, J.M. Polo, A.R.R. Forrest, A.K. Nowak, I. Voineagu, L. Martelotto, and R. Lister. 2022. Human prefrontal cortex gene regulatory dynamics from gestation to adulthood at single-cell resolution. Cell. 185:4428-4447.e4428.

Huang, Y., K.E. Happonen, P.G. Burrola, C. O’Connor, N. Hah, L. Huang, A. Nimmerjahn, and G. Lemke. 2021. Microglia use TAM receptors to detect and engulf amyloid β plaques. Nature Immunology. 22:586–594.

Inuzuka, H., H. Fukushima, S. Shaik, P. Liu, A.W. Lau, and W. Wei. 2011. Mcl-1 ubiquitination and destruction. Oncotarget. 2:239–244.

Jayadev, S., A. Case, B. Alajajian, A.J. Eastman, T. Möller, and G.A. Garden. 2013. Presenilin 2 influences miR146 level and activity in microglia. Journal of neurochemistry. 127:592–599.

Jayadev, S., A. Case, A.J. Eastman, H. Nguyen, J. Pollak, J.C. Wiley, T. Möller, R.S. Morrison, and G.A. Garden. 2010. Presenilin 2 Is the Predominant γ-Secretase in Microglia and Modulates Cytokine Release. PloS one. 5.

Kaja, S., N. Sumien, V.V. Shah, I. Puthawala, A.N. Maynard, N. Khullar, A.J. Payne, M.J. Forster, and P. Koulen. 2015. Loss of spatial memory, learning and motor coordination during normal aging is accompanied by changes in brain presenilin 1 and 2 expression levels. Molecular Neurobiology. 52:545–554.

Kang, J., and J. Shen. 2020. Cell-autonomous role of Presenilin in age-dependent survival of cortical interneurons. Molecular neurodegeneration. 15:72.

Kelley, K.W., H. Nakao-Inoue, A.V. Molofsky, and M.C. Oldham. 2018. Variation among intact tissue samples reveals the core transcriptional features of human CNS cell classes. Nature neuroscience. 21:1171–1184.

Kemmerling, N., P. Wunderlich, S. Theil, B. Linnartz-Gerlach, N. Hersch, B. Hoffmann, M.T. Heneka, B.d. Strooper, H. Neumann, and J. Walter. 2017. Intramembranous processing by γ-secretase regulates reverse signaling of ephrin-B2 in migration of microglia. Glia. 65:1103–1118.

Kim, S., S.H. Cho, K.Y. Kim, K.Y. Shin, H.S. Kim, C.H. Park, K.A. Chang, S.H. Lee, D. Cho, and Y.H. Suh. 2009. Alpha-synuclein induces migration of BV-2 microglial cells by up-regulation of CD44 and MT1-MMP. Journal of neurochemistry. 109:1483–1496.

Kimberly, W.T., M.J. LaVoie, B.L. Ostaszewski, W. Ye, M.S. Wolfe, and D.J. Selkoe. 2003. γ-Secretase is a membrane protein complex comprised of presenilin, nicastrin, aph-1, and pen-2. Proceedings of the National Academy of Sciences of the United States of America. 100:6382–6387.

Kimberly, W.T., W. Xia, T. Rahmati, M.S. Wolfe, and D.J. Selkoe. 2000. The transmembrane aspartates in presenilin 1 and 2 are obligatory for γ-secretase activity and amyloid β-protein generation. The Journal of biological chemistry. 275:3173–3178.

Kopan, R., and M.X. Ilagan. 2009. The canonical Notch signaling pathway: unfolding the activation mechanism. Cell. 137:216–233.

Lauer, A.A., J. Mett, D. Janitschke, A. Thiel, C.P. Stahlmann, C.M. Bachmann, F. Ritzmann, B. Schrul, U.C. Müller, R. Stein, M. Riemenschneider, H.S. Grimm, T. Hartmann, and M.O.W. Grimm. 2020. Regulatory feedback cycle of the insulin-degrading enzyme and the amyloid precursor protein intracellular domain: Implications for Alzheimer’s disease. Aging cell. 19:e13264.

Lee, M.K., H.H. Slunt, L.J. Martin, G. Thinakaran, G. Kim, S.E. Gandy, M. Seeger, E. Koo, D.L. Price, and S.S. Sisodia. 1996. Expression of presenilin 1 and 2 (PS1 and PS2) in human and murine tissues. The Journal of Neuroscience. 16:7513–7525.

Lessard, C.B., E. Rodriguez, T.B. Ladd, L.M. Minter, B.A. Osborne, L. Miele, T.E. Golde, and Y. Ran. 2019. Individual and combined presenilin 1 and 2 knockouts reveal that both have highly overlapping functions in HEK293T cells. The Journal of biological chemistry. 294:11276–11285.

Luo, W.J., H. Wang, H. Li, B.S. Kim, S. Shah, H.J. Lee, G. Thinakaran, T.W. Kim, G. Yu, and H. Xu. 2003. PEN-2 and APH-1 coordinately regulate proteolytic processing of presenilin 1. The Journal of biological chemistry. 278:7850–7854.

Manocha, G.D., A.M. Floden, K. Rausch, J.A. Kulas, B.A. McGregor, L. Rojanathammanee, K.R. Puig, K.L. Puig, S. Karki, M.R. Nichols, D.C. Darland, J.E. Porter, and C.K. Combs. 2016. APP Regulates Microglial Phenotype in a Mouse Model of Alzheimer’s Disease. The Journal of Neuroscience. 36:8471–8486.

Mastrangelo, P., P.M. Mathews, M.A. Chishti, S.D. Schmidt, Y. Gu, J. Yang, M.J. Mazzella, J. Coomaraswamy, P. Horne, B. Strome, H. Pelly, G. Levesque, C. Ebeling, Y. Jiang, R.A. Nixon, R. Rozmahel, P.E. Fraser, P. St George-Hyslop, G.A. Carlson, D. Westaway, and S.B. Prusiner. 2005. Dissociated Phenotypes in Presenilin Transgenic Mice Define Functionally Distinct γ-Secretases. Proceedings of the National Academy of Sciences of the United States of America. 102:8972–8977.

Meckler, X., and F. Checler. 2016. Presenilin 1 and presenilin 2 target γ-secretase complexes to distinct cellular compartments. The Journal of biological chemistry. 291:12821–12837.

Muller, P.R., T.J. Lee, W. Zhi, S. Kumar, S. Vyavahare, A. Sharma, V. Kumar, C.M. Isales, M. Hunter, and S. Fulzele. 2023. Proteomic Analysis of Female Synovial Fluid to Identify Novel Biomarkers for Osteoarthritis. Life. 13:605.

Peng, W., Y. Xie, C. Liao, Y. Bai, H. Wang, and C. Li. 2022. Spatiotemporal patterns of gliosis and neuroinflammation in presenilin 1/2 conditional double knockout mice. Front Aging Neurosci. 14.

Pfaffl, M.W. 2001. A new mathematical model for relative quantification in real-time RT-PCR. Nucleic Acids Res. 29:e45–e45.

Pimenova, A.A., and A.M. Goate. 2020. Novel presenilin 1 and 2 double knock-out cell line for in vitro validation of PSEN1 and PSEN2 mutations. Neurobiology of disease:104785.

Placanica, L., L. Tarassishin, G. Yang, E. Peethumnongsin, S.-H. Kim, H. Zheng, S.S. Sisodia, and Y.-M. Li. 2009a. Pen2 and presenilin-1 modulate the dynamic equilibrium of presenilin-1 and presenilin-2 γ-secretase complexes. The Journal of biological chemistry. 284:2967–2977.

Placanica, L., L. Zhu, and Y.-M. Li. 2009b. Gender- and age-dependent gamma-secretase activity in mouse brain and its implication in sporadic Alzheimer disease. PloS one. 4:e5088–e5088.

Ran, F.A., P.D. Hsu, J. Wright, V. Agarwala, D.A. Scott, and F. Zhang. 2013. Genome engineering using the CRISPR-Cas9 system. Nature Protocols. 8:2281.

Rangaraju, S., E.B. Dammer, S.A. Raza, P. Rathakrishnan, H. Xiao, T. Gao, D.M. Duong, M.W. Pennington, J.J. Lah, N.T. Seyfried, and A.I. Levey. 2018. Identification and therapeutic modulation of a pro-inflammatory subset of disease-associated-microglia in Alzheimer’s disease. Molecular neurodegeneration. 13:24.

Ross, R.A., J.D. Walton, D. Han, H.F. Guo, and N.K. Cheung. 2015. A distinct gene expression signature characterizes human neuroblastoma cancer stem cells. Stem cell research. 15:419–426.

Ryman, D.C., N. Acosta-Baena, P.S. Aisen, T. Bird, A. Danek, N.C. Fox, A. Goate, P. Frommelt, B. Ghetti, J.B.S. Langbaum, F. Lopera, R. Martins, C.L. Masters, R.P. Mayeux, E. McDade, S. Moreno, E.M. Reiman, J.M. Ringman, S. Salloway, P.R. Schofield, R. Sperling, P.N. Tariot, C. Xiong, J.C. Morris, R.J. Bateman, and N. and the Dominantly Inherited Alzheimer. 2014. Symptom onset in autosomal dominant Alzheimer disease: A systematic review and meta-analysis. Neurology. 83:253–260.

Sannerud, R., C. Esselens, P. Ejsmont, R. Mattera, L. Rochin, Arun K. Tharkeshwar, G. De Baets, V. De Wever, R. Habets, V. Baert, W. Vermeire, C. Michiels, Arjan J. Groot, R. Wouters, K. Dillen, K. Vints, P. Baatsen, S. Munck, R. Derua, E. Waelkens, Guriqbal S. Basi, M. Mercken, M. Vooijs, M. Bollen, J. Schymkowitz, F. Rousseau, Juan S. Bonifacino, G. Van Niel, B. De Strooper, and W. Annaert. 2016. Restricted location of PSEN2/γ-secretase determines substrate specificity and generates an intracellular Aβ pool. Cell. 166:193–208.

Saura, C.A., S.-Y. Choi, V. Beglopoulos, S. Malkani, D. Zhang, B.S.S. Rao, S. Chattarji, R.J. Kelleher Iii, E.R. Kandel, K. Duff, A. Kirkwood, and J. Shen. 2004. Loss of Presenilin Function Causes Impairments of Memory and Synaptic Plasticity Followed by Age-Dependent Neurodegeneration. Neuron. 42:23–36.

Shen, J., R.T. Bronson, D.F. Chen, W. Xia, D.J. Selkoe, and S. Tonegawa. 1997. Skeletal and CNS Defects in Presenilin-1-Deficient Mice. Cell. 89:629–639.

Soto-Faguás, C.M., P. Sanchez-Molina, and C.A. Saura. 2021. Loss of presenilin function enhances tau phosphorylation and aggregation in mice. Acta neuropathologica communications. 9:162.

Steiner, H., E. Winkler, D. Edbauer, S. Prokop, G. Basset, A. Yamasaki, M. Kostka, and C. Haass. 2002. PEN-2 is an integral component of the γ-secretase complex required for coordinated expression of presenilin and nicastrin. The Journal of biological chemistry. 277:39062–39065.

Tang, N., and K.P. Kepp. 2018. Aβ42/Aβ40 Ratios of Presenilin 1 Mutations Correlate with Clinical Onset of Alzheimer’s Disease. Journal of Alzheimer’s Disease. 66:939–945.

Thakur, M.K., and S. Ghosh. 2007. Age and sex dependent alteration in presenilin expression in mouse cerebral cortex. Cellular and molecular neurobiology. 27:1059–1067.

Thinakaran, G., C.L. Harris, T. Ratovitski, F. Davenport, H.H. Slunt, D.L. Price, D.R. Borchelt, and S.S. Sisodia. 1997. Evidence that levels of presenilins (PS1 and PS2) are coordinately regulated by competition for limiting cellular factors. The Journal of biological chemistry. 272:28415–28422.

Watanabe, H., K. Imaizumi, T. Cai, Z. Zhou, T. Tomita, and H. Okano. 2021. Flexible and accurate substrate processing with distinct presenilin/γ-secretases in human cortical neurons. eNeuro. 8:ENEURO.0500-0520.2021.

Watanabe, H., M. Iqbal, J. Zheng, M. Wines-Samuelson, and J. Shen. 2014. Partial Loss of Presenilin Impairs Age-Dependent Neuronal Survival in the Cerebral Cortex. The Journal of Neuroscience. 34:15912–15922.

Wines-Samuelson, M., E.C. Schulte, M.J. Smith, C. Aoki, X. Liu, R.J. Kelleher, III, and J. Shen. 2010. Characterization of Age-Dependent and Progressive Cortical Neuronal Degeneration in Presenilin Conditional Mutant Mice. PloS one. 5:e10195.

Wong, P.C., H. Zheng, H. Chen, M.W. Becher, D.J.S. Sirinathsinghji, M.E. Trumbauer, H.Y. Chen, D.L. Price, L.H.T. Van Der Ploeg, and S.S. Sisodia. 1997. Presenilin 1 is required for Notch 1 and DII 1 expression in the paraxial mesoderm. Nature. 387:288–292.

Wunderlich, P., K. Glebov, N. Kemmerling, N.T. Tien, H. Neumann, and J. Walter. 2013. Sequential proteolytic processing of the triggering receptor expressed on myeloid cells-2 (TREM2) protein by ectodomain shedding and γ-secretase-dependent intramembranous cleavage. The Journal of biological chemistry. 288:33027–33036.

Xia, W., P. Bringmann, J. McClary, P.P. Jones, W. Manzana, Y. Zhu, S. Wang, Y. Liu, S. Harvey, M.R. Madlansacay, K. McLean, M.P. Rosser, J. MacRobbie, C.L. Olsen, and R.R. Cobb. 2006. High levels of protein expression using different mammalian CMV promoters in several cell lines. Protein Expression and Purification. 45:115–124.

Yang, D.-S., A. Tandon, F. Chen, G. Yu, H. Yu, S. Arawaka, H. Hasegawa, M. Duthie, S.D. Schmidt, T.V. Ramabhadran, R.A. Nixon, P.M. Mathews, S.E. Gandy, H.T.J. Mount, P. St George-Hyslop, and P.E. Fraser. 2002. Mature glycosylation and trafficking of nicastrin modulate its binding to presenilins. The Journal of biological chemistry. 277:28135–28142.

Yonemura, Y., E. Futai, S. Yagishita, C. Kaether, and S. Ishiura. 2016. Specific combinations of presenilins and Aph1s affect the substrate specificity and activity of γ-secretase. Biochemical and biophysical research communications. 478:1751–1757.

Yonemura, Y., E. Futai, S. Yagishita, S. Suo, T. Tomita, T. Iwatsubo, and S. Ishiura. 2011. Comparison of presenilin 1 and presenilin 2 γ-secretase activities using a yeast reconstitution system. The Journal of biological chemistry. 286:44569–44575.

Yu, H., C.A. Saura, S.-Y. Choi, L.D. Sun, X. Yang, M. Handler, T. Kawarabayashi, L. Younkin, B. Fedeles, M.A. Wilson, S. Younkin, E.R. Kandel, A. Kirkwood, and J. Shen. 2001. APP Processing and Synaptic Plasticity in Presenilin-1 Conditional Knockout Mice. Neuron. 31:713–726.

Zhao, Y., C.Y. Zeng, X.H. Li, T.T. Yang, X. Kuang, and J.R. Du. 2020. Klotho overexpression improves amyloid-β clearance and cognition in the APP/PS1 mouse model of Alzheimer’s disease. Aging cell. 19:e13239.

Zhu, M., F. Gu, J. Shi, J. Hu, Y. Hu, and Z. Zhao. 2008. Increased oxidative stress and astrogliosis responses in conditional double-knockout mice of Alzheimer-like presenilin-1 and presenilin-2. Free Radical Biology and Medicine. 45:1493–1499.

