## Supplementary material for "Differential expression and modulation of presenilin-1 and presenilin-2 in neural cell lines": Eccles - Differential expression and modulation of presenilin-1 and presenilin-2 in neural cell lines - SI Material

Correspondence to: **Melissa K. Eccles**

Edinburgh EH4 2XU, UK

Key Words: Presenilin-1, Presenilin-2, Alzheimer’s disease, neurons, microglia

Funding Information: MKE - Dementia Australia | Dementia Australia Research Foundation (DARF)

Abbreviations: Aβ, amyloid-β; AD, Alzheimer's disease; CNS, central nervous system; PS-Std, presenilin fusion standard; PS1, presenilin-1; PS2, presenilin-2; PS1γ, PS1-γ-secretase; PS2γ, PS2-γ-secretase.

Running Title: Differential expression of presenilin proteins

### SUPPORTING INFORMATION

#### *SI Table 1: CRISPR-Cas9 gRNA target sequences for PS1 and PS2 knockout*

| Cell Line | Target Gene | gRNA site 1 | gRNA site 2 |
|---|---|---|---|
| HMC3 | PSEN1 | $^{5'}$GTTTCAACCAGCATACGAAG$^{3'}$ | $^{5'}$TAAAACCTATAACGTTGCTG$^{3'}$ |
| | PSEN2 | $^{5'}$GCTCCCCTACGACCCGGAGA$^{3'}$ | $^{5'}$ACGATCATGCACAGAGTGAC$^{3'}$ |
| M17 | PSEN1 | $^{5'}$TTATCTAATGGACGACCCCA$^{3'}$ | $^{5'}$GAGCAATACTGTACGTAGCC$^{3'}$ |
| | PSEN2 | $^{5'}$GCTCCCCTACGACCCGGAGA$^{3'}$ | $^{5'}$ACGATCATGCACAGAGTGAC$^{3'}$ |

#### *SI Table 2: qPCR primer sequences*

| Gene Target | Forward Primer | Reverse Primer |
|---|---|---|
| PSEN1 | $^{5'}$CCAGAGGAAAGGGGAGTAAAACTT$^{3'}$ | $^{5'}$ACAGGCTATGGTTGTGTTCCA$^{3'}$ |
| PSEN2 | $^{5'}$TCATCTGCCATGGTGTGGAC$^{3'}$ | $^{5'}$GTCTTCTTCCATCTCCGGGT$^{3'}$ |
| APH1a | $^{5'}$GGTGTTTTTCGGCTGCACTT$^{3'}$ | $^{5'}$CAGAAAAATGCCCCTGCGAC$^{3'}$ |
| APH1b | $^{5'}$CTGCGCCTTCATTGCCTTC$^{3'}$ | $^{5'}$GAAGAAAGCTCCGGCGATGA$^{3'}$ |
| NCSTN | $^{5'}$ACTAGCAGGTTTGTGCAGGG$^{3'}$ | $^{5'}$TCTGATGAGTGGCGTTGAGC$^{3'}$ |
| PSENEN | $^{5'}$TGCCTTTTCTCTGGTTGGTCA$^{3'}$ | $^{5'}$CGCCAGACATAGCCTTTGAT$^{3'}$ |
| TMEM119¶ | $^{5'}$CTTCCTGGATGGGATAGTGGAC$^{3'}$ | $^{5'}$GCACAGACGATGAACATCAGC$^{3'}$ |
| P2RY12 | $^{5'}$CCACTCTGCAGGTTGCAATAAC$^{3'}$ | $^{5'}$TTGCATTTCTTGTTGGTTACCTGA$^{3'}$ |
| RPLP0¶ | $^{5'}$GAAACTCTGCATTCTCGCTTCC$^{3'}$ | $^{5'}$GAAACTCTGCATTCTCGCTTCC$^{3'}$ |
| CDC73 | $^{5'}$CCTGGCGTCGTGATTAGTGAT$^{3'}$ | $^{5'}$TCGAGCAAGACGTTCAGTCC$^{3'}$ |
| CD200 | $^{5'}$GTGCACAGCACAAGTGCAAG$^{3'}$ | $^{5'}$TGGGCATTTTGCAGAGAGCA$^{3'}$ |
| CX3CL1 | $^{5'}$CCACGGTGTGACGAAATGC$^{3'}$ | $^{5'}$GTCTCGTCTCCAAGATGATTGC$^{3'}$ |
| POLR2A | $^{5'}$TCCTCGCATGATTGTCACCC$^{3'}$ | $^{5'}$GTTCATCACTTCACCCCGCT$^{3'}$ |
| GAPDH§ | $^{5'}$CTGCTTTTAACTCTGGTAAAGT$^{3'}$ | $^{5'}$GCGCCAGCATCGCCCCA$^{3'}$ |

¶ TMEM119 and RPLP0 primers are from Bennet et al. (2016)

§ GAPDH primers are from Koch et al. (2012)


Bennett, M. L., Bennett, F. C., Liddelow, S. A., Ajami, B., Zamanian, J. L., Fernhoff, N. B., Mulinyawe, S. B., Bohlen, C. J., Adil, A., Tucker, A., Weissman, I. L., Chang, E. F., Li, G., Grant, G. A., Hayden Gephart, M. G., and Barres, B. A. (2016) New tools for studying microglia in the mouse and human CNS. Proceedings of the National Academy of Sciences 113, E1738-E1746

Koch, P., Tamboli, I. Y., Mertens, J., Wunderlich, P., Ladewig, J., Stüber, K., Esselmann, H., Wiltfang, J., Brüstle, O., and Walter, J. (2012) Presenilin-1 L166P mutant human pluripotent stem cell-derived neurons exhibit partial loss of γ-secretase activity in endogenous amyloid-β generation. The American Journal of Pathology 180, 2404-2416

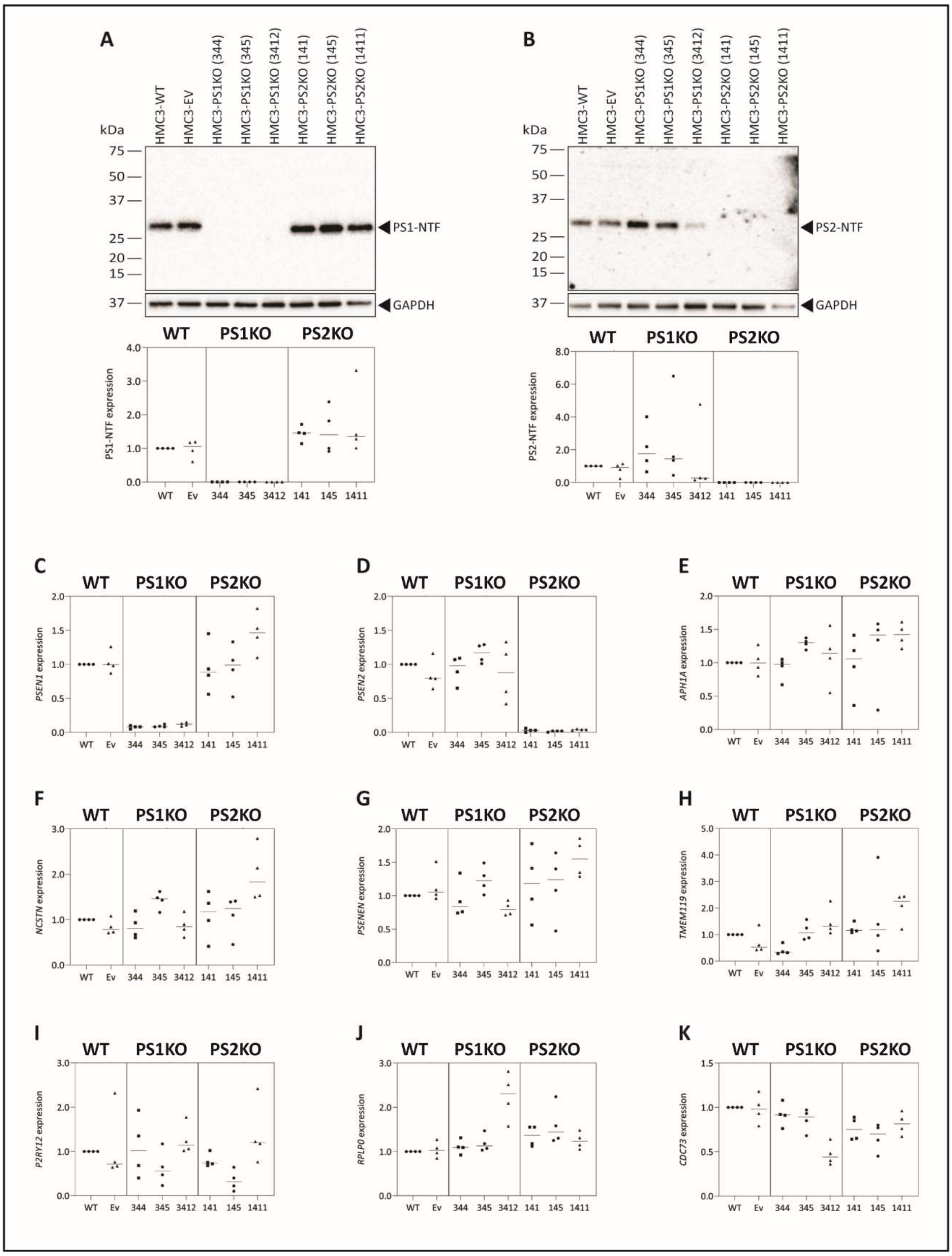


***SI Figure 1: Characterisation of HMC3 PS1KO and PS2KO clones***

Assessment of HMC3 PS1KO and PS2KO clones via immunbloting of PS1-NTF (A), PS2-NTF (B) and qPCR of PSEN1 (C), PSEN2 (D), APH1A (E), NCSTN (F), PSENEN (G), TMEM119 (H), P2RY12 (I), RPLP0 (J), CDC73 (K).

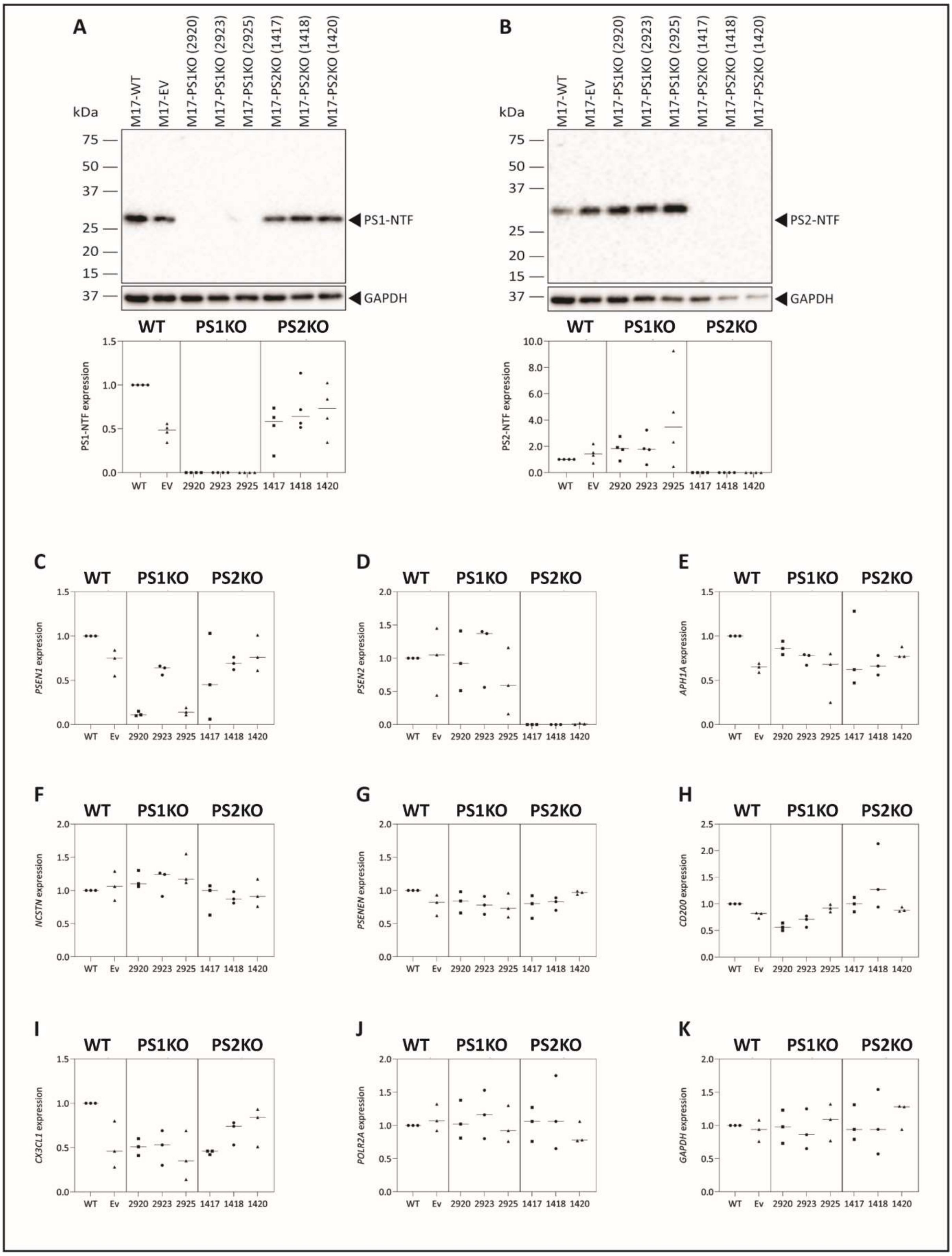


***SI Figure 2: Characterisation of M17 PS1KO and PS2KO clones***

Assessment of M17 PS1KO and PS2KO clones via immunbloting of PS1-NTF (A), PS2-NTF (B) and qPCR of PSEN1 (C), PSEN2 (D), APH1A (E), NCSTN (F), PSENEN (G), CD200 (H), CX3CL1 (I), POLR2A (J), GAPDH (K).
